# EoRNA, a barley gene and transcript abundance database

**DOI:** 10.1101/2020.11.24.395749

**Authors:** Linda Milne, Micha Bayer, Paulo Rapazote-Flores, Claus-Dieter Mayer, Robbie Waugh, Craig G Simpson

## Abstract

A high-quality, barley gene reference transcript dataset (BaRTv1.0), was used to quantify gene and transcript abundances from 22 RNA-seq experiments, covering 843 separate samples. Using the abundance data we developed a Barley Expression Database (EoRNA* – Expression of RNA) to underpin a visualisation tool that displays comparative gene and transcript abundance data on demand as transcripts per million (TPM) across all samples and all the genes. EoRNA provides gene and transcript models for all of the transcripts contained in BaRTV1.0, and these can be conveniently identified through either BaRT or HORVU gene names, or by direct BLAST of query sequences. Browsing the quantification data reveals cultivar, tissue and condition specific gene expression and shows changes in the proportions of individual transcripts that have arisen via alternative splicing. TPM values can be easily extracted to allow users to determine the statistical significance of observed transcript abundance variation among samples or perform meta analyses on multiple RNA-seq experiments. * Eòrna is the Scottish Gaelic word for Barley

## Background & Summary

Barley is one our earliest domesticated crops and is used for food and processed as malt to produce beer and spirits. It is a widely studied crop model with abundant genetic resources that include diverse natural cultivated, wild and landrace collections, experimentally constructed populations, introgression and mutant lines. Its robust diploid genetics are supported by numerous high-resolution linkage maps and fully sequenced reference and pan-genome sequences (1, 2, 3, 4, 5). Genomic diversity has contributed to barley being grown worldwide, producing harvestable yields under a broad range of environmental conditions and climates (1, 4, 6). As a direct consequence, variation in gene expression contributes implicitly to its adaptive response. Plant gene expression constantly changes throughout the day, throughout plant development and responds to changing environmental conditions, providing a mechanism for different genotypes to react and adapt to both transient and chronic stresses (For example, 7, 8, 9, 10, 11, 12, 13).

Although the responses of individual genes to specific genetic, biological or environmental interventions are frequently described, whole transcriptome responses over multiple growth stages and conditions, and consequently the network of genes and transcripts involved in these responses, are largely unknown. As growth, morphology and physiology vary substantially among barley genotypes, either when individual genotypes are grown under different conditions or when different genotypes are grown under identical conditions, their transcriptomes reveal a landscape that is highly dynamic, adaptable and unique to the applied conditions (**14, 15)**. This is not simply the product of the regulation of gene expression at the level of transcription. Differentially abundant precursor messenger RNAs (pre-mRNAs) may be further subjected to alternative splice site selection, forming an assembly of specific transcript isoforms. (12, 13, 16, 17, 18). The cellular transcriptome is therefore comprised of transcripts derived from a combination of both transcriptional and post-transcriptional processes.

A high confidence barley reference transcript dataset (BaRTv1.0) represented by 60,444 gene models and 177,240 transcript sequences are provided in a database (https://ics.hutton.ac.uk/barleyrtd/index.html) that positions the transcripts on the barley cv. Morex reference genome version 1 (19). The database is fully searchable using either BaRT or HORVU gene names from the Barley cv Morex pseudomolecules, by key word annotation or by BLAST sequence searches. The database provides best BLAST homologies of the longest transcript to Arabidopsis, rice and Brachypodium, and provides links to GO annotations and GO enrichment studies. The BaRTv1.0 reference transcript dataset (RTD) enables rapid and precise quantification using non-alignment bioinformatic tools such as Kallisto and Salmon from short-read RNA-seq data (20, 21). Levels of expression from these tools are measured in Transcripts per million (TPM) for a given BaRTv1.0 transcript (22).

In summary, to highlight the utility of the barley RTD coupled to transcript quantification with Salmon, we quantified gene and transcript abundances from 22 separate RNA-seq studies, covering 843 samples from a broad range of different tissues, conditions and genotypes. Our aim was to allow rapid and intuitive access to the transcript quantification values of each of these RNA-seq studies without considering any experimental batch, sample or study variation and without making any statement about significant changes in gene expression across the different studies. We make the resource available to the community via the EoRNA database web site (https://ics.hutton.ac.uk/eorna/index.html) to simplify and accelerate exploration of the abundance of target transcripts from individual or groups of genes. The numerical TPM data can be downloaded for further expression analysis or for meta-analysis of barley RNA-seq datasets to support investigations into transcriptional responses among tissues/organs or as a result of different interventions, allowing the identification of genes and transcripts commonly expressed across multiple studies (23, 24, 25, 26). Intuitive transcript abundance plots graphically illustrate tissue and condition specific gene expression and alternative splicing.

## Methods

### Selected RNA-seq datasets and data processing

A total of 22 publicly available RNA-seq datasets consisting of 843 samples including replicates were downloaded from NCBI - Sequence Read Archive database (https://www.ncbi.nlm.nih.gov/sra/) to quantify against the barley RTD (BaRTv1.0) (Supplementary Table S1). All datasets were produced using Illumina platforms and were selected with mostly >90 bp and paired-end reads with a quality of q >=20. All raw data were processed using Trimmomatic-0.30 (27) using default settings to preserve a minimum Phred score of Q20 over 60 bp. One of the samples (NOD1) was over-represented with respect to read numbers due to a repeat run being necessary and was therefore subsampled to 60 million reads. Read quality checks before and after trimming were performed using FastQC (fastqc_v0.11.5) (https://www.bioinformatics.babraham.ac.uk/projects/fastqc/).

### Generation of the EoRNA database

A database and website front-end were constructed to allow easy access to BaRTv1.0 transcripts and expression analyses using the LAMP configuration (Linux, Apache, mySQL, and Perl). Additional annotation was added to the transcripts by homology searching against the predicted peptides from rice (rice pseudo-peptides v 6.0; (28)) and from Arabidopsis thaliana (TAIR pseudo-peptides v 10, The Arabidopsis Information Resource) using BLASTX at an e-value cutoff of less than 1e-50 (29). The website https://ics.hutton.ac.uk/eorna/index.html allows users to interrogate data through an entry point via three methods: (i) a BLAST search of the reference barley assembly or the predicted transcripts; (ii) a keyword search of the derived rice and Arabidopsis thaliana BLAST annotation, and; (iii) a direct string search using the transcript, gene, or contig identifiers. To distinguish this set of predicted genes and transcripts from previously published ‘MLOC_’ and HORVU identifiers, genes were prefixed as ‘BART1_0-u00000’ for the unpadded or ‘BART1_0-p00000’ for the padded QUASI version, with BART1_0-p00000.000 representing the individual transcript number. The RNA-seq TPM values are shown in interactive stacked bar plots produced with plotly R libraries (https://plotly.com/r/) and the TPM values are also available as a text file for each gene. The exon structures of the transcripts for each gene are shown in graphical form, and links to the transcripts themselves provides access to the transcript sequences in FASTA format. Each transcript has also been compared to the published set of predicted genes (HORVUs) to provide backwards compatibility.

### GO annotation

Transcript sequences were translated to protein sequences using TransDecoder (https://github.com/TransDecoder/TransDecoder/wiki). Gene Ontology (GO) annotation was then determined by running all 60,444 genes in BaRTv1.0 through Protein ANNotation with Z-score (PANNZER) (**30.** Koskinen et al., 2015). GO annotations were based on predicted proteins with ORF >100 amino acids and orthologues found in the Uniprot database. Output annotations were placed in a lookup table with text descriptions about protein functionality.

## Data Records

BaRTv1.0 and BaRTv1.0 – QUASI are available as .fasta and .GFF files and can be downloaded from https://ics.hutton.ac.uk/barleyrtd-new/downloads.html. An additional version of the RTD is available in the Zenodo repository (http://doi.org/10.5281/zenodo.3360434).

The results matrix containing all the TPM values across all 843 samples for all 177,240 BaRTv1.0 transcripts can be downloaded directly along with the metadata file from https://ics.hutton.ac.uk/eorna/download.html. An additional version of the results matrix and metadata file is available in the Zenodo repository (http://doi.org/10.5281/zenodo.4286079). To develop the plots and create the transcript abundance values (TPMs) publicly available sequences from the Sequence Read Archive (SRA) or European Nucleotide Archive (ENA) were used (accession numbers: PRJEB13621; PRJEB18276; PRJNA324116; PRJEB12540; PRJEB8748; PRJNA275710; PRJNA430281; PRJNA378582; PRJNA378723; PRJNA439267; PRJNA396950; PRJDB4754; PRJNA428086; PRJEB21740; PRJEB25969; PRJNA378334; PRJNA315041; PRJNA294716; PRJEB14349; PRJEB32063; PRJEB19243; PRJNA558196. Metadata on these datasets can be found in Supplementary Tables 1 and 2.

## Technical Validation

### BaRTv1.0 database and expression plots

The BaRTv1.0 reference transcript dataset consists of 60,444 genes and 177,240 transcripts mapped to the cv. Morex pseudomolecules. To access the barley reference transcript dataset a public database and website front-end were constructed to allow researchers to download the reference transcript dataset and interrogate the data via a BLAST search, keyword search or string search using the BaRT or HORVU gene/transcript identifiers (https://ics.hutton.ac.uk/barleyrtd/index.html) (19). The transcripts are arranged as gene models and viewed through GBrowse. Transcript sequences are given in FASTA format and homologies of the longest transcripts are compared to Arabidopsis, Rice and Brachypodium. Until now, Salmon calculated TPM values for each gene across 16 different tissues/developmental stages in both graphic and tabular formats is presented. Since the initial publication, the BaRTV1.0 database has continued to evolve and we have established Gene Ontology (GO) annotation for 26,794 genes using Protein ANNotation with Z-score (PANNZER) (**30.** Koskinen et al., 2015) with text descriptions about protein functionality and provided a lookup table for download.

### EoRNA database - Quantification of multiple RNA-seq samples and expression plots

Establishing BaRTv1.0 has facilitated the precise quantification of RNA transcript abundance from any barley short-read RNA-seq dataset. We used BaRTv1.0 to quantify transcript abundance and diversity observed in a collection of 22 Illumina short-read RNA-seq experiments, 18 of which were obtained from the short-read archive (SRA) and the remainder produced in-house. Each RNA-seq experiment was given a label that contained the letter E (referring to external datasets) followed by a number or the letter I (internal datasets) followed by a number. The datasets contained a total of 843 samples and 3,762 Gbp of expressed sequences. They come from both barley landraces and cultivars, an array of organs and tissues at different developmental stages, and plants/seedlings grown under a range of biotic and abiotic stresses (Supplementary Table S1 and S2). Most RNA-seq datasets consisted of paired-end reads (90 - 150 bp in length) and were produced using Illumina HiSeq 2000, 2500, 4000 or HiSeq X instruments. Exceptions were the dataset from Golden Promise anthers and meiocytes, which contained over 2 billion paired end 35-76 bp reads. The raw RNA-seq data from all samples was trimmed and adapters removed using Trimmomatic and quality controlled using FastQC. TPM values were calculated individually for all 843 RNA-seq samples using Salmon (version Salmon-0.8.2) using BaRTv1.0-QUASI, a ‘padded’ version of BaRTv1.0 which has been shown to improve transcript quantification, as the reference transcript dataset (**19.** Rapazote-Flores et al., 2019). As BaRTv1.0 was assembled using the cv. Morex reference genome, we first assessed the mapping rates from all samples, including those from other genotypes. The Morex samples showed an average mapping rate of 94.39% (SD 8.18%) while the remaining samples, which consisted of 60 different barley genotypes showed a slightly reduced mapping rate of 92.32% (SD 4.93%) (Supplementary Table 3).

Salmon estimates the relative abundance of different transcript isoforms in the form of transcripts per million (TPM), a commonly used normalization method computed using the library size, number of reads and the effective length of the transcript. (20, 21). The EoRNA data provides an opportunity to examine the effect of the normalisation procedure across many diverse samples. Regression analyses was used to explore the raw read counts and different versions of normalised counts by library size and effective length of the transcript. Good normalisation procedures will remove most of the dependency on these variables such that the output of regression analysis represented by the R-square value (which measures the percentage of variation accounted for) can be used to compare different normalisations. Here, an R-square value closer to zero indicates effective normalisation. For efficient calculation, we first reduced the number of transcripts by selecting those which had non-zero values in at least 80% of the samples. This left 32739 transcripts over the 843 samples and gave 27,598,977 values to study how different normalisation approaches accounted for variation between experiments. Regression analysis was used first to explore the relationship between raw read counts by library size and length of the transcript, which gave an adjusted R-squared value of 1.28% indicating low predictive value within the dataset. Transposing variables to a log-scale increased the R-square to 10.68%, which suggested a far stronger predictive value on this scale and shows that a large amount of variation in the raw counts can be removed by log-transforming. Replacing the log counts with normalised data using Salmon’s effective transcript length, which corrects for transcript length bias (20), reduced the adjusted R-square value to 0.09%. This compared to normalisation by RPKM (Reads Per Kilobase per Million and normalizes the raw read count by transcript length and sequencing depth) (adjusted R-square of 0.57%) or TPMs calculated by transcript length alone (adjusted R-square of 0.62%). In summary, the normalised TPM outputs from Salmon using an effective transcript length reduced variability such that most of the dependency on library size and transcript length was removed (Supplementary data 1; Supplementary Table 4).

The normalised output TPM values from Salmon were collated and plotted using plotly R libraries (https://plotly.com/r/) to allow quick subjective and interactive comparisons in transcript abundance levels between the samples. The TPM values for each gene/plot are also given as a text file for download. We chose to plot the graphs as the TPM values without log scaling, to show the additive changes between the samples and replicates.

### Expression plot utility

Stacked bar graph plots display the TPM values calculated by Salmon for all 60,444 genes in the database for all 843 samples, representing over 50 million plot points. The x-axis displays the 843 samples versus the y-axis which displays transcript abundance in each sample as TPM values (Figure 1). Each individual sample bar graph stacks the TPM values contributed by each gene transcript to permit simple visualisation of the differences in transcript abundances between different samples and helps identify the predominant transcript(s) for that gene. Each plot may be scanned interactively to activate a label that gives information on the RNA-seq experiment, sample run number, tissue and treatment for that sample (from the metadata table, Supplementary Table 2). Users can zoom in to focus on individual experiment and sample plots. Without processing the data or assigning any statistical significance to the graphs, the results presented allow the researcher to determine whether their gene(s) of interest are expressed in the different experiments and among samples within an experiment. Large changes in TPM abundances were observed between the samples for many genes. For example, BaRT1_u-31819 showed altered gene expression in the root meristematic zone compared to the root elongation and maturation zones in the E1 dataset, which is further supported by expression in the root tissue in the I1 dataset (Figure 1).

**Figure 1.**
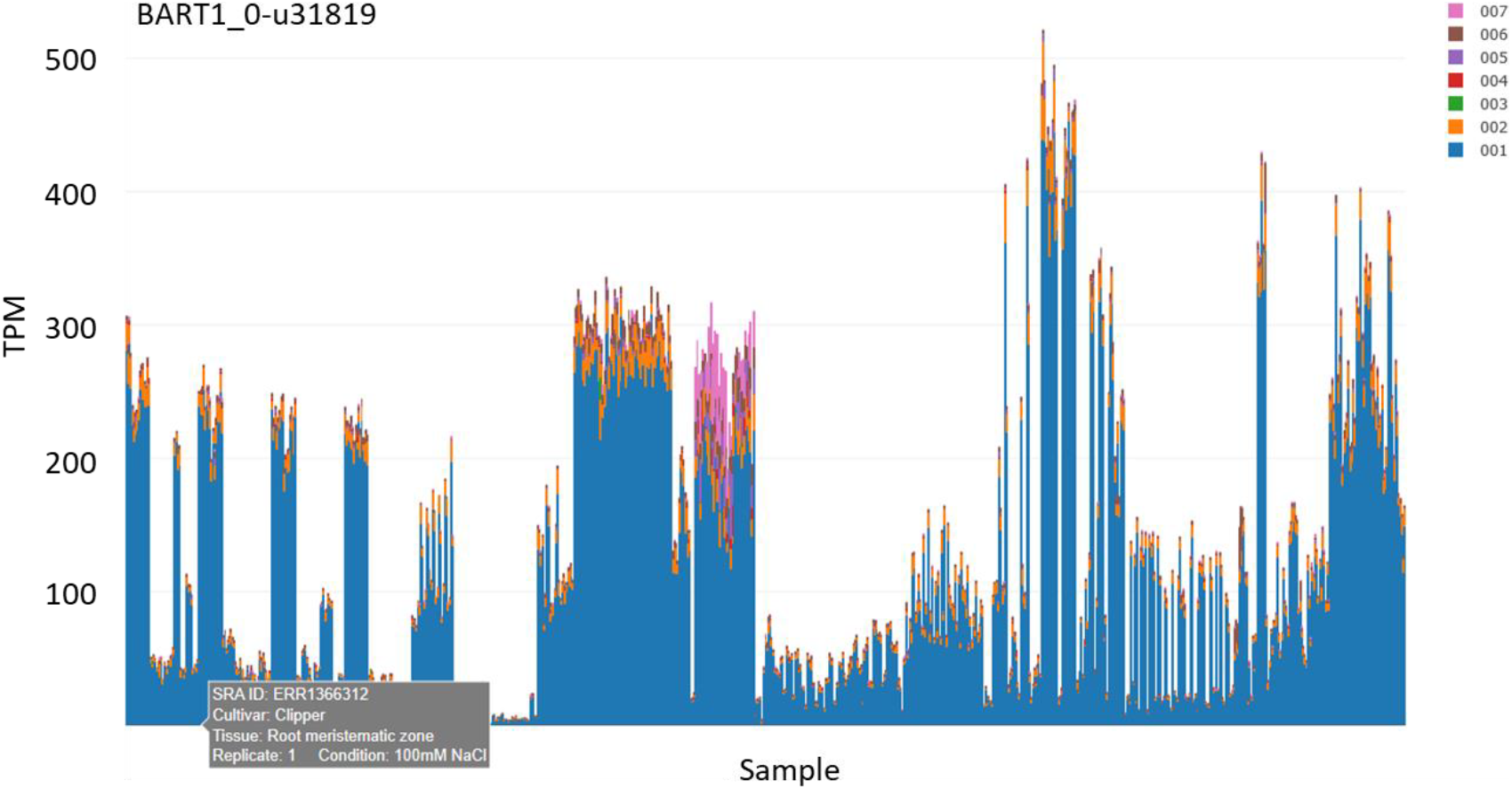
Variable expression between RNA-seq samples. The plot represents transcript abundances as transcripts per million (TPM) across 843 samples for BaRT1_0-u31919 (similarity to a small nuclear ribonucleoprotein family protein). Different colours represent different transcripts for that gene. Scanning over the plot gives a label describing cultivar, tissue, experimental condition (if available), replicate number and the short-read archive sequencing read number.

### Tissue specific expression

The experimental panel of 22 RNA-seq datasets were from a broad range of cultivars, tissues, organs and biotic and abiotic conditions. The interactive plots enable the user to quickly identify potential candidate genes that show a high degree of tissue specificity. For example, BART1_0-u49225 (with similarity to a UDP-Glycosyltransferase superfamily protein) was specifically and highly expressed to over 1,000 TPM in developing grain 15 days post anthesis (I1) and in developing barley spikes that contain developing grain (E20). Expression was segregating in hulless barley grain in recombinant inbred lines that were used to assess glucan content (E10). (Figure 2A). BART1_0-u14427 was highly abundant only in tissues subjected to low temperature stress (E2 and I2) (Figure 2B) and BART1_0-u50915 is one of a number of barley Pathogenesis-related 1 protein genes that was induced to over 10,000 TPM in response to Cochliobolus sativus (E19) and Fusarium graminearum (E20) (Figure 2C).

**Figure 2.**
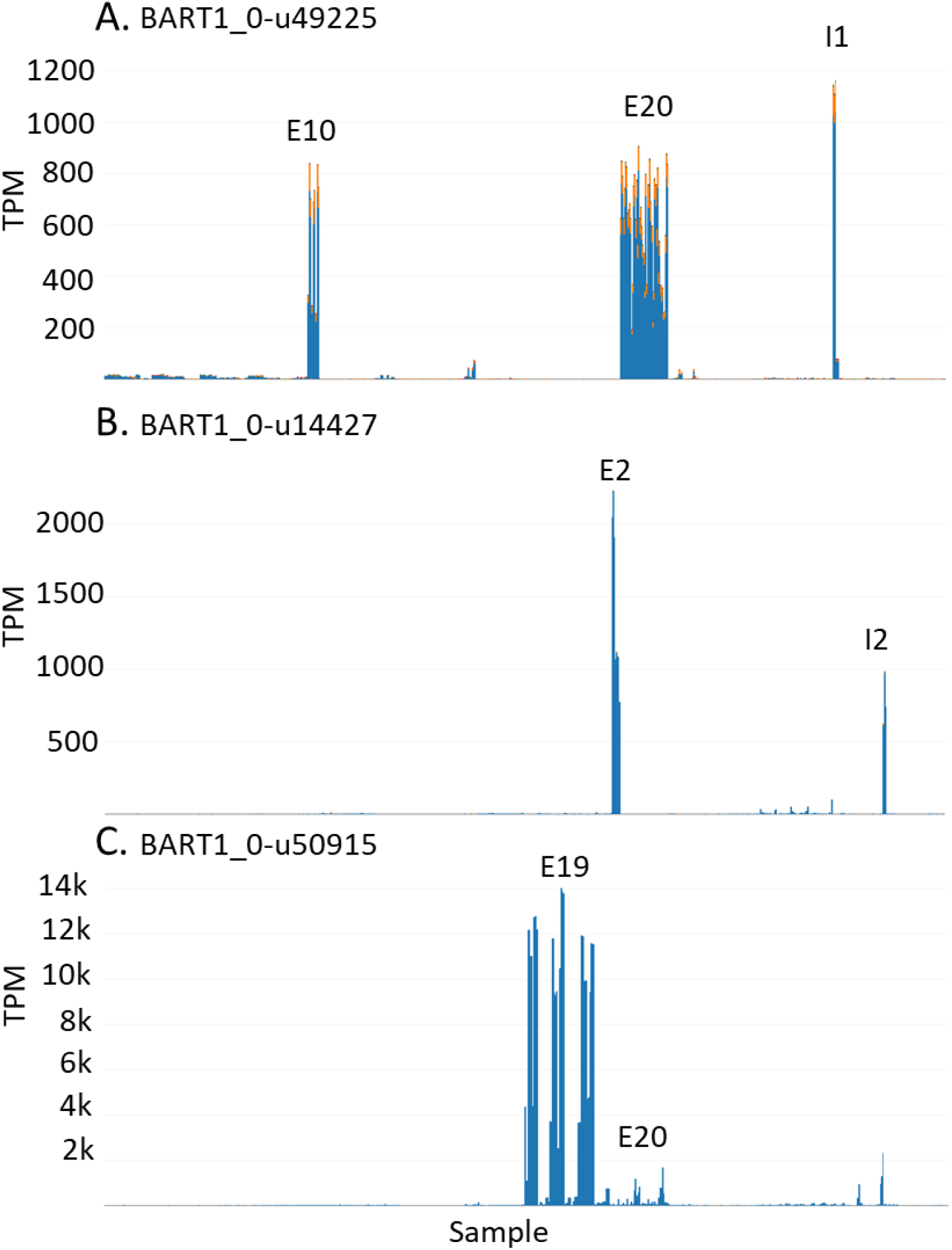
Tissue and condition specific expression. A. BART1_0-u49225 specific expression in developing grain tissue used in experimental RNA-seq datasets E10, E20 and I1. B. BART1_0-u14427 specific expression in low temperature stress RNA-seq datasets E2 and I2. C. BART1_0-u50915 specific expression in response to pathogen RNA-seq datasets E19 and E20.

### Confirmatory expression

Interactive plots may be used to investigate the expression of genes that have been previously studied in a limited number of tissues/cultivars or using a different expression platform and consequently expands expression analysis across the range of tissues that are currently in EoRNA. For example, we previously described the expression of INTERMEDIUM-C (BART1_0-u26546; HORVU4Hr1G007040), a modifier of lateral spikelet fertility in barley and an ortholog of the maize domestication gene TEOSINTE BRANCHED 1. Microarray analysis of 15 tissues showed that transcript abundance was low with greatest expression in the developing inflorescence (31). The RNA-seq panel here confirmed low abundances for this gene across all the samples (<7.5 TPM), with greatest expression in shoot apices (E7); apical meristems (E13) and developing spikes at the awn primordium stage (E14) (Figure 3).

**Figure 3.**
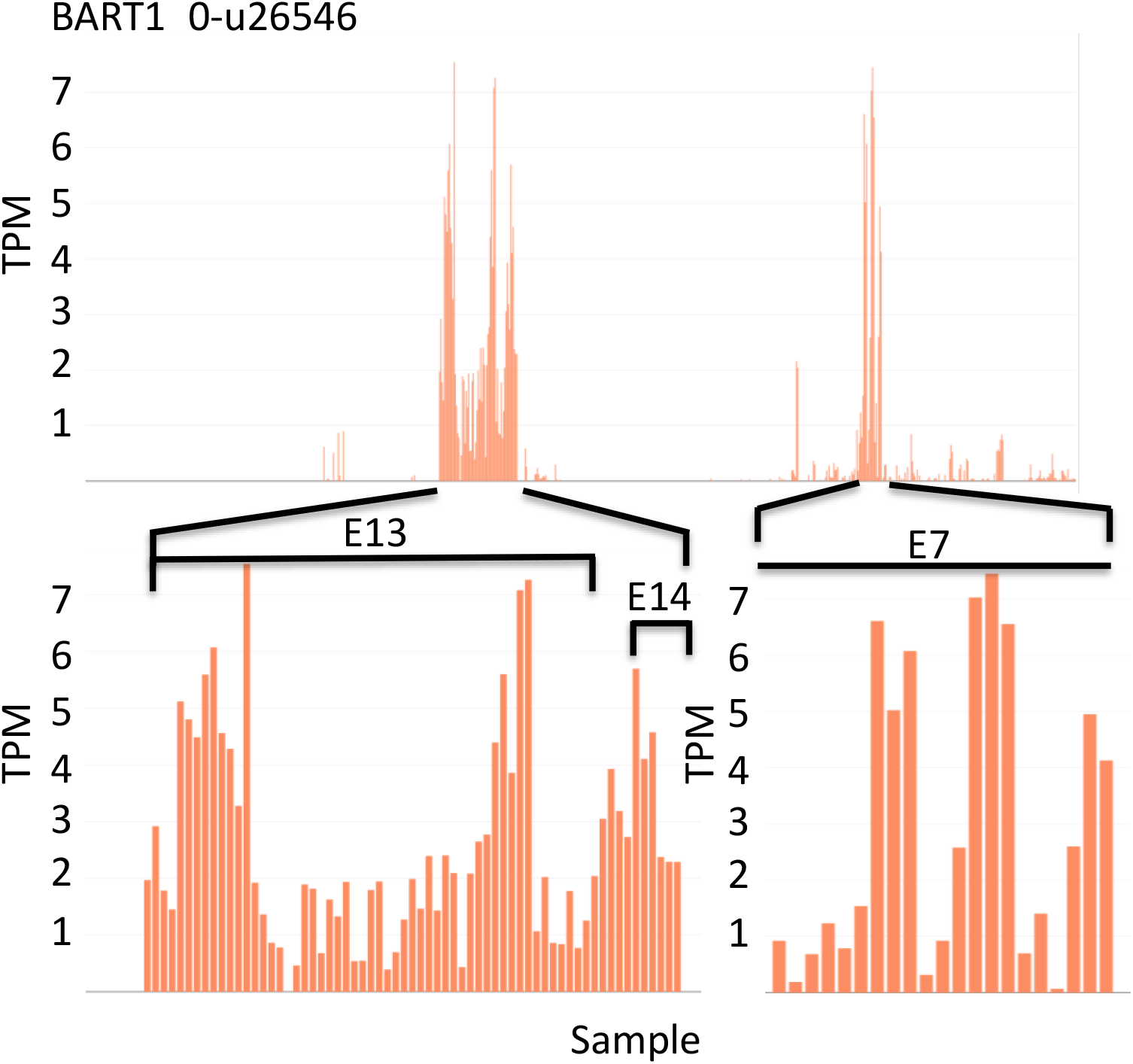
Abundance levels of INTERMEDIUM-C (HvTB1) (BART1_0-u26546) across the 22 RNA-seq experiments. E7 – Photoperiod response RNA-seq dataset from shoot apex; E13 - Six Rowed - VRS3 RNA-seq dataset from apical meristems; E14 - Floret development RNA-seq dataset from developing spikes at awn primordium stage. Abundances given in Transcripts per million (TPM). The bottom Panel shows zoomed-in regional views.

### Segregation expression

The RNA-seq datasets consist of several experiments that contain mutant lines targeted to specific genes, recombinant inbred lines (RILs) and near isogenic lines (NILs). The expression of genes found at quantitative trait loci, or through genome-wide association studies show changes in gene expression at these loci between the parents and in the population. The seed longevity experiment (E17) illustrated gene expression changes in RILs and NILs from the landraces L94 (short-lived seeds) and Cebada capa (long-lived seeds). QTL analysis identified three QTLs on 1H (SLQ1.1 to 1.3) and a single QTL on 2H (SLQ2). Gene expression analysis identified differentially expressed genes positioned within the SLQ1 and 2 regions (32). Using the interactive plots confirmed the barley population expression pattern of these differentially expressed genes. The plots show changes among the parental types retained in the recombinant inbred and near isogenic lines (Figure 4). For example, BART1_0-u01011(MLOC_61374) is positioned within SLQ1.1 and showed low expression in Cebada capa and the NILs at SLQ1.1 (Figure 4A) and BART1_0-u15865 (MLOC_73587) showed expression in Cebada capa that was absent in L94 and found expressed in SLQ2 NILs Figure 4B). The transcript abundances of these genes were shown in the context of the remaining 21 RNA-seq experiments tested.

**Figure 4.**
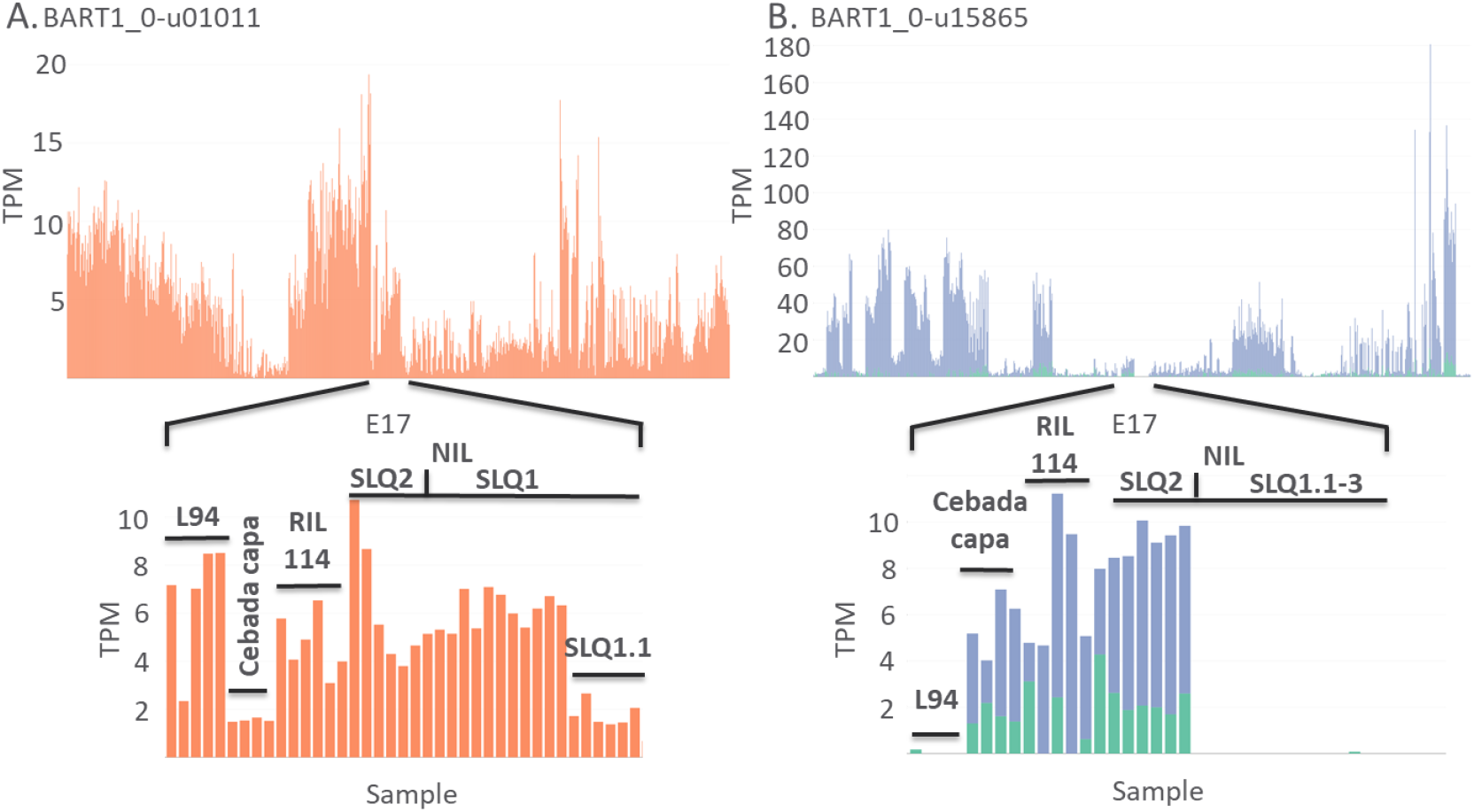
Abundance levels of differentially expressed genes at quantitative trait loci. Detailed abundances (TPM) are shown for a seed longevity experiment (E17) between parents (L94 and Cebada capa), recombinant inbred lines (RIL114) and near isogenic lines to the L94 parent and showing variation at QTLs SLQ1 and SLQ1-3. A. BART1_0-u01011(MLOC_61374) is located at SLQ1.1 and B. BART1_0-u15865 (MLOC_73587) is located at SLQ2.

### Gene targeted mutations

Unless a mutation either specifically impacts sequences governing the expression of a target gene, or removes all or part of a gene by deletion, then the outcome of a mutation on observed transcript abundance may vary substantially, resulting in loss, reduced, maintained or increased transcript levels. The interactive plots allow researchers to observe rapidly and intuitively the effect of a mutation on the expression of a target gene and, based on the experimental design, the possible trans-acting effects on the expression of other genes. For example, experiment E19 consists of a series of disease resistance tests on cv. Morex and a gamma irradiation induced Morex mutant (14-40) selected for its susceptibility to spot blotch (Bipolaris sorokiniana). The expression of BART1_0-u18601; HORVU3Hr1G019920 (glycine-rich protein) and BART1_0-u41161; HORVU5Hr1G120850 (similarity to a long-chain-fatty-acid—CoA ligase 1) were knocked out in the mutant, which is clearly observed in the interactive plots (33) (Figure 5).

**Figure 5.**
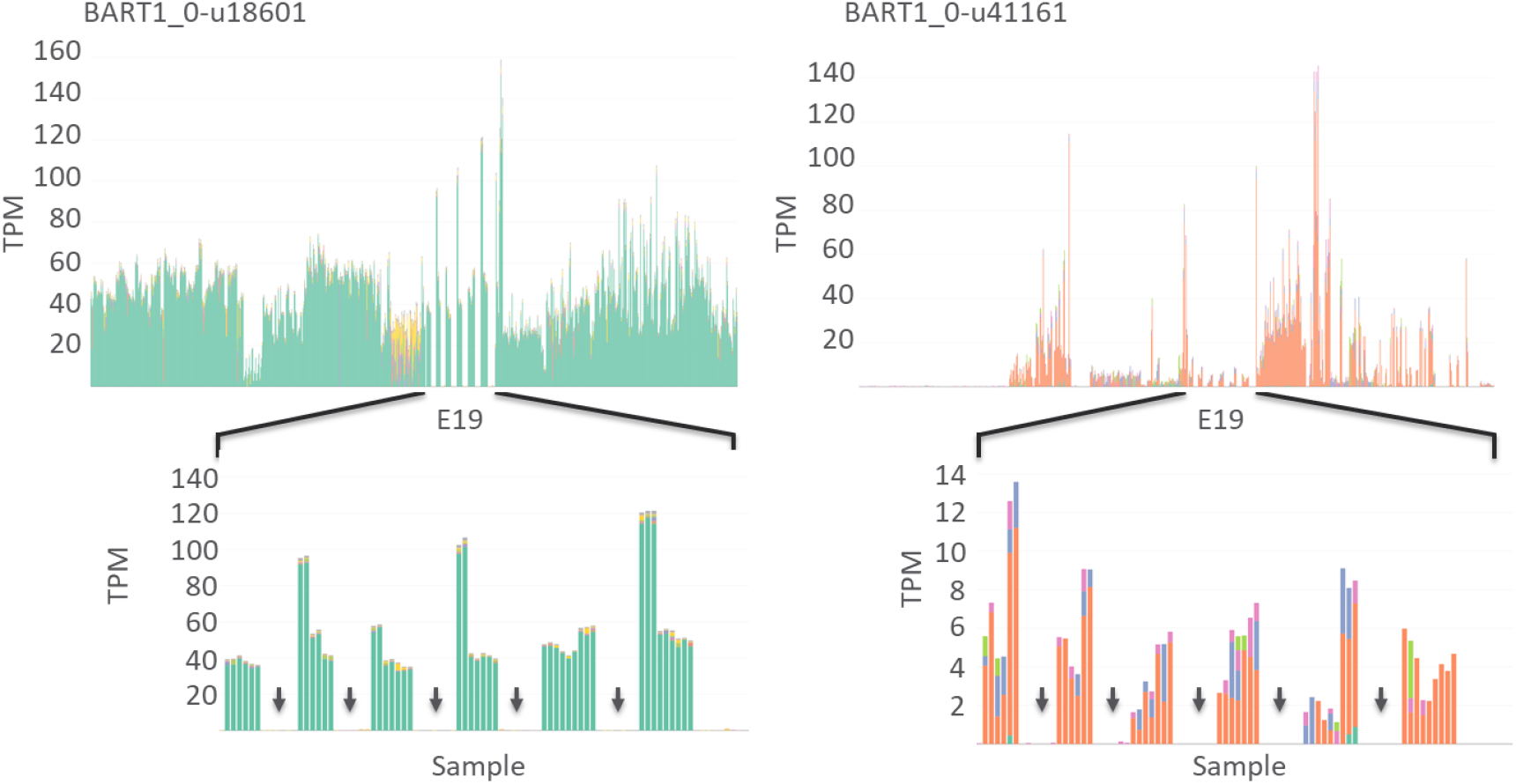
Expression knockout in a mutant background. The pattern of transcript abundances of two genes (BART1_0-u18601 and BART1_0-u4116) is shown across all the samples and given in Transcripts per million (TPM). Detailed transcript abundances are shown for the E19 RNA-seq dataset - RNA-seq of Hordeum vulgare inoculated with Cochliobolus sativus. The gaps arrowed between the expression in the wild type cv. Morex are multiple samples derived from the barley cv. Morex mutant 14-40, which shows disruption of expression.

### Transcript variation between cultivars

To create the BaRTv1.0 RTD, transcripts from multiple datasets from a range of tissues, treatments and cultivars were mapped to cv. Morex pseudomolecules to maximise read coverage support for genes and splice junctions (19). BaRTv1.0 is, therefore, a predominantly cv. Morex RTD. Nevertheless, transcripts that contain indels in other cultivars will be found in BaRTv1.0. Salmon quantifications of the 843 individual samples was able to identify and quantify cultivar specific transcripts. BaRT1_u-06868 showed a selection of different transcripts due to genotype differences. Alignment with genomic sequence and the most highly abundant transcripts shows a small run of 4 GCAG repeats in one genotype compared to a run of 3 GCAG repeats in a different genotype. These genotype specific variant transcripts were observed across the range of cultivars used in the RNA-seq experiments. For example, the experimental dataset E1 shows two different cultivars cvs. Clipper and Sahara with two different main transcript variants, which is the result of the 4bp indel. Clipper shows use of the transcripts .001 and .002 while Sahara uses transcripts .005 and .006 (Figure 6). The transcriptome assemblies and quantifications using BaRTv1.0 shows that cultivar specific transcripts can be easily distinguished.

**Figure 6.**
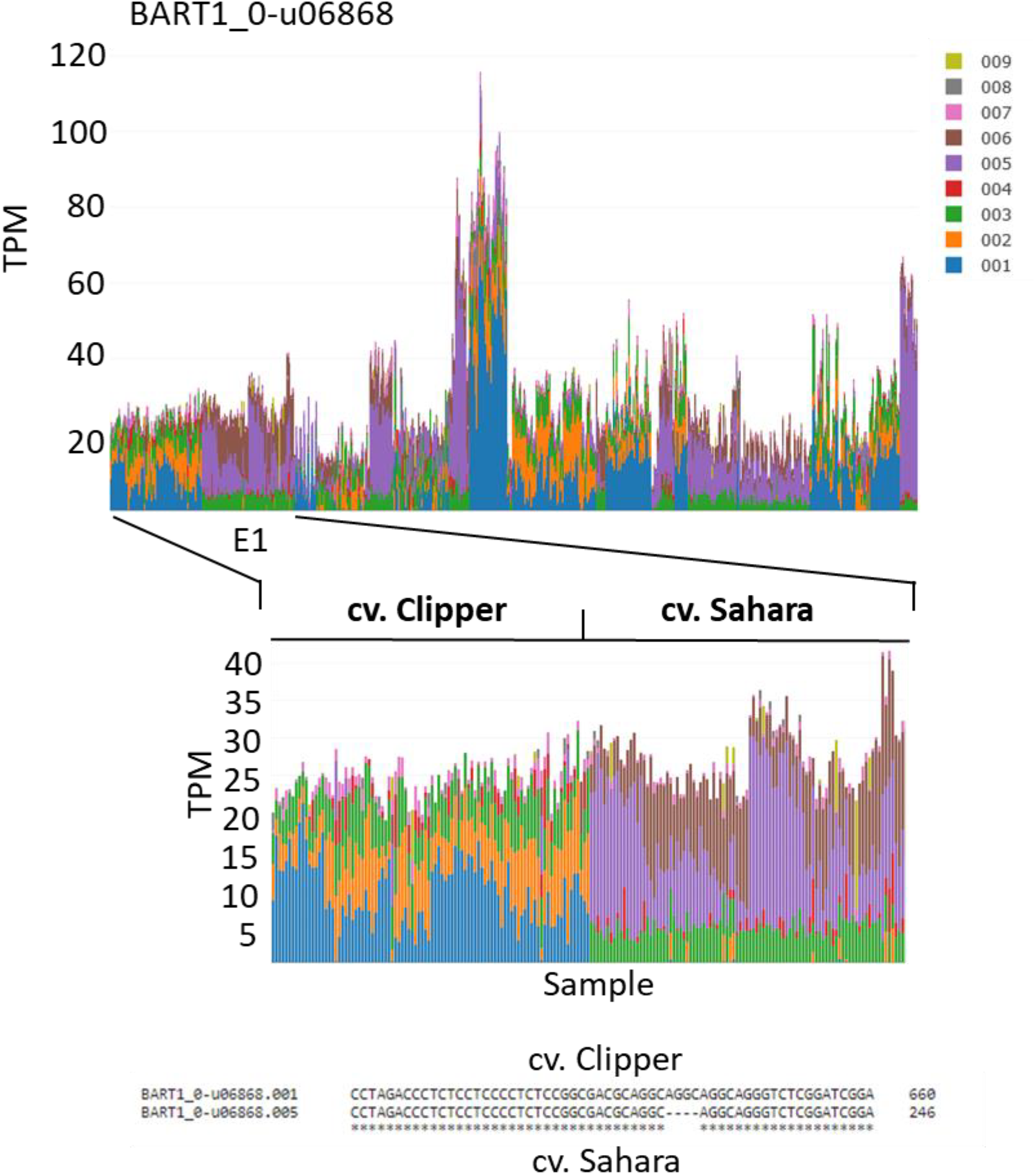
Transcripts that represent allelic variants across barley cultivars. BaRT1_u-06868 shows transcripts .001 (blue) and .002 (orange in the cv. Clipper, while cv. Sahara shows transcripts .005 (purple) and .006 (brown). Sequence alignment between transcripts .001 and .005 shows the 4bp deletion in cv. Sahara found in transcript 005.

#### Alternative splice site switching

Selection of alternative splice sites results in the formation of multiple alternative transcripts. The proportions of alternative transcripts may change in different tissues or as the result of a changing environment. Many of these changes require detailed analysis to determine significant changes in the amounts and proportions of the alternative transcripts. Nevertheless, the stacked bar graphs allow large changes in the abundance of alternative transcripts to be detected between samples. For example, BaRT1_u-00022 was expressed across all tissues but in some samples an alternative transcript, BaRT1_u-00022.001, shown in blue, predominated over BaRT1_u-00022.003 shown in green (Figure 7A). The difference between the two transcripts was an alternative intron in the 3’UTR, which was retained in transcript .001 and spliced out in transcript .003. Comparison with the meta-data (Supplementary table 2) showed tissue specific abundance of transcript .001 in grain/caryopsis and germinating grain (coleoptiles) in the experimental datasets E8, E10, E17, I1 and I2. Comparison across the different experiments and replicates supports both the tissue and cultivar specific variation. For example, the alternative .001 transcript was also observed in Golden Promise in datasets E11 and I6. The plots also illustrate dynamic changes in alternative splicing in different tissues or because of different stresses. For example, BaRT1_u-40919, which has similarity to a cold inducible Zinc finger-containing glycine-rich RNA-binding protein, shows switching of transcript .001 to .005 during cold stress, which is the result of the selection of an alternative intron (I2) (Figure 7B). In both these cases, the reading frame of the protein is unaffected but extends the length of the 3’UTR in the transcripts where the intron is retained. These examples highlight transcript variation because of genotypic differences and dynamic alternative splicing as a result of tissue/organ specific splicing or changing environmental conditions.

**Figure 7.**
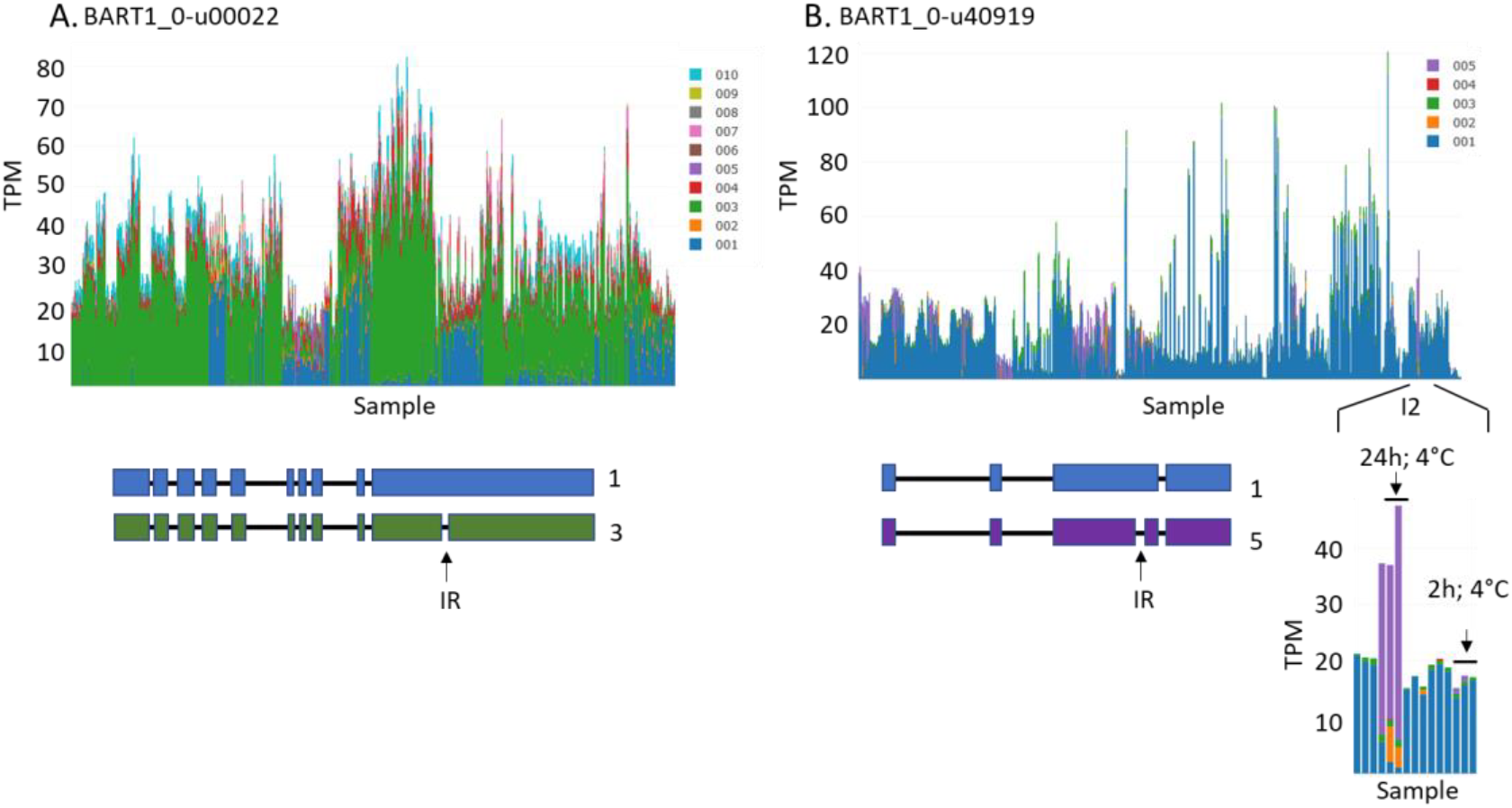
Alternative transcripts across the RNA-seq experiments. Different colours on the stacked bar graph indicate different gene transcripts produced through alternative splicing. Expression levels given in TPM – transcripts per million. A. BaRT1_u-00022 shows two main transcripts in blue (.001) and green (.003). B. BaRT1_u-40919 shows transcript switching in the cold response experimental set I2. Alternative splicing leads to switching from transcript .001 (blue) to .005 (purple in the cold. Gene models for each gene are presented and the position of the retained intron (IR) shown.

## Discussion

Comprehensive reference transcript datasets are required for rapid, accurate quantification of gene expression using RNA-seq. Quantification at the transcript level further allows robust and routine analysis of alternative splicing (34, 35, 36). Here we used the barley reference transcript dataset, BaRTv1.0, to demonstrate the value and utility of a barley RTD for gene expression studies and AS analysis. We used BaRTv1.0 to quantify transcripts in 22 RNA-seq datasets, covering multiple genotypes, tissues and different abiotic and biotic stress conditions. BaRTv1.0 was assembled against the cv. Morex genome, but in this analysis we used RNA-seq data from a wide-range of cultivars and lines and found that mapping rates in all cultivars remained high (94.39% in cv. Morex compared to 92.32% in the other cultivars). We found expression and alternative splicing abundances varied between cultivars, tissues/organs and between environmental changes and stresses. The data is presented in a single accessible database that gives visual and numerical access to expression data for barley genes across all the tested barley samples (https://ics.hutton.ac.uk/eorna/index.html).

The importance of comparing between sample sets allows researchers to answer how their gene of interest is expressed in other tissues or under what condition. RNA-seq expression results are displayed in graphical form, simply as TPM values directly from the outputs of Salmon, without considering batch differences that may occur between samples, among experimental studies and does not assign statistical significances. We recognise that to include statistical analysis and thereby define significant DE or DAS would require complete control over experimental design, sample preparation and sequencing analysis. These interactive plots, therefore, simply permit rapid visual assessment of expression levels of selected genes of interest. TPM values are accessible and allow users to perform their own DE and DAS analysis, such as found in the 3D RNA-seq interactive graphical user interface (37) or by comparing multiple RNA-seq datasets by meta-analysis methods (23, 24, 25, 26). The results will enable the construction of transcript/co-expression/regulatory networks and support the development of proteomic resources for barley.

We did not carry out validation experiments using alternative methods, such as RT-PCR, as we do not have access to all the RNA samples used to produce the RNA-seq data. However, multiple RNA-seq samples consisted of similar tissues or conditions that showed similar gene expression responses. This was particularly noticeable in the genes that showed tissue or condition specific expression, such as those from developing grain tissue, low temperature stress and in response to pathogens (Figure 2). In addition, we have previously performed RT-PCR alternative splicing validation experiments on 5 of the tissues in the I1 RNA-seq experiment and found a strong correlation (r2=0.83) with the alternatively spliced transcript proportions of RNA-seq, supporting the ability of the RNA-seq data to accurately detect changes in AS (19).

Output expression values such as TPM from RNA-seq experiments are under continuous discussion and development and may be affected by sequencing protocols and experimental conditions (38). Here, TPM values were calculated using Salmon to allow transcript abundances to be compared between samples. To check that the TPM values were representative as expression values, we determined variability across all the samples using linear regression analyses and found that the output from Salmon showed the lowest variability and therefore provided the best normalisation across all the samples. Some of the downloaded RNA-seq datasets revealed experimental samples that had extremely low or high read depths and poor mapping rates that after normalisation suggested abnormally high TPM values. These were not included in our analyses (data not shown).

We have given examples of genes that clearly illustrate the wide utility offered by access to datasets from multiple RNA-seq experiments. The plots identified genes that were uniquely expressed in a cultivar, tissue or condition specific manner. Considering the range of samples displayed, the unique abundances in specific samples support the potential value of these genes as expression ‘biomarkers’ for that tissue or condition. There were other uniquely expressed genes found in the interactive plots and only three were reported here to illustrate utility:-BART1_0-u49225, with similarity to a UDP-Glycosyltransferase family member, was found specifically expressed in developing grain; BART1_0-u14427, with similarity to late embryogenesis abundant (LEA) proteins was induced after 24 h at low temperature; and BART1_0-u50915, which is one of a number of barley Pathogenesis-related 1 protein genes that are established pathogen responsive genes (Figure 2). The plots also identified cis- and trans-acting induced expression (or loss of expression) of genes that segregate among near isogenic lines or mutant populations (Figure 4 and 5) and cultivar specific transcripts (Figure 6). The expression characteristics may help identify, retain or exclude candidate genes from involvement in a given biological process, form the basis for the development of tissue specific reporter genes, validate observed expression QTL or explore the genomic landscape of actively expressed genes.

Barley exhibits a high frequency of alternative splicing that impacts development and adaptation to the surrounding daily and seasonal environment. The plots revealed genes that change their splice site selection patterns in different tissues and organs and, in some cases, show switching in splice site selection as a response to stress (Figure 7). In addition, genotypic differences in diverse barley cultivars and landraces can lead to considerable changes in the gene expression. Single nucleotide polymorphisms or insertion/deletions at important splice sites and in splicing regulatory elements can affect the abundance of transcript isoforms and alter translational reading frames or transcript stability. An example here shows how a 4 bp deletion in cv. Sahara led to selection of two different transcripts in the BaRT RTD by cv. Clipper. The functional impact of genetic variations on splicing diversity will impact phenotypic diversity and cultivar adaptation to local environments.

BaRT is under constant incremental improvement. The next release of BaRT is being developed by incorporating new short and, importantly, long-read RNA-seq datasets. The need to capture the diversity of different transcripts from a wider range of genotypes will further lead to the development of a pan-transcriptome barley RTD to match a barley pan-genome sequence (5, 39). This will ultimately result in recalculation of the TPM values. In addition, new RNA-seq experiments are constantly submitted to the sequence archives. We are currently developing a pipeline that allows automated addition of newly deposited RNA-seq datasets associated with subsequent quantification using the latest RTD and updated releases of EoRNA. This will continually expand the utility of the interactive plots and provide straightforward and open access of RNA-seq data to researchers, adding considerable value to the stand-alone RNA-seq datasets. In summary, the BaRT RTD is part of a unique pipeline that facilitates fast robust routine quantification of barley gene transcripts, visualised in EoRNA through interactive plots linked to gene models and metadata, ultimately leading to robust and consistent estimation of barley gene expression and alternative splicing across multiple samples.

## Usage Notes

The expression data is most easily accessible through an intuitive and easy to use Web interface: https://ics.hutton.ac.uk/eorna/index.html.

Gene and transcript sequence information and expression data can be accessed through Homology Searches, Annotation Searches or thorough BLAST nucleotide or protein sequences. Barley Pseudomolecule gene names (HORVU numbers) can be easily translated to BART identifiers.

The plots showing individual gene expression across all the samples has a link under the plot to a text delimited file with all the expression (TPMs), tissue, condition, cultivar and replicate. The whole dataset describing expression of all the BaRT genes can downloaded as a single txt delimited file. This is further stored at http://doi.org/10.5281/zenodo.4286079.

## Acknowledgements

The authors wish to acknowledge critical reading by Peter E. Hedley at The James Hutton Institute. This research was supported and developed by Scottish Government Rural and Environment Science and Analytical Services division (RESAS) and funding from the Biotechnology and Biological Sciences Research Council (BBSRC) (BB/I00663X/1: A draft sequence of the barley genome) and ERC project 669182 ‘SHUFFLE’ to RW.

## Author contributions

PR-F and MB downloaded and assembled the RNA-seq datasets. LM established the searchable database. LM, MB, CS, and C-DM conceived and designed the interactive plots for the database. PR-F, MB, LM, CS, and C-DM performed the analysis of the RNA-seq data and outputs. CS, LM, MB, C-DM and RW wrote the paper.

